# Integrating structural homology with deep learning to achieve highly accurate protein-protein interface prediction for the human interactome

**DOI:** 10.1101/2025.06.09.658393

**Authors:** Dapeng Xiong, Mateo Torres, Diana Murray, Zizhao Zhang, Le Li, Aniket C. Naravane, Robert Fragoza, Barry Honig, Haiyuan Yu

**Author notes:** These authors contributed equally to this work (Dapeng Xiong, Mateo Torres). Corresponding authors (B.H.) and (H.Y.).

## Abstract

A significant portion of disease-causing mutations occur at protein-protein interfaces however, the number of structurally resolved multi-protein complexes is extremely small. Here we present a computational pipeline, PIONEER2, that integrates 3D structural similarity with geometric deep learning to accurately predict protein binding partner-specific interfacial residues. We compare the performance of PIONEER2 to that of AlphaFold3 and found, using a test set of PDB structures, that their performance is quite similar. However, about 20% of AlphaFold3 predictions for protein-protein complexes in the PDB have AlphaFold3 ranking scores below 0.5, which indicates an uncertain model. For these structures, PIONEER2 outperforms AlphaFold3 at discriminating interfacial from non-interfacial residues. Further, about half of the AlphaFold3 ranking scores on high confidence protein-protein interactions (PPIs) not associated with a PDB structure are below 0.5 indicating that PIONEER2 offers superior interface prediction for a large number of PPIs for which structures are not available. We created a comprehensive 3D structurally informed interactome encompassing all 352,124 experimentally detected binary human PPIs in the current literature and made PIONEER2 interface predictions for each. We experimentally validated these predictions by generating 1,866 mutations and testing their disruptive impact on 5,010 mutation-interaction pairs. PIONEER2-predicted interfaces are found to be comparable to PDB structures in their ability to predict disruptive mutations while AlphaFold3 performance is reduced. Similarly, PIONEER2-predicted interfaces outperform AlphaFold3 in accounting for the depletion of non-deleterious common population variants and the enrichment of disease-related mutations on protein surfaces. Overall, our results suggest that PIONEER2-predicted interfaces provide a valuable tool for studying disease etiology, advancing personalized medicine and for fundamental research. We further implemented PIONEER2 as a user-friendly web server (https://pioneer2.yulab.org) platform for users to explore our 3D interactome models and conduct genome-wide functional genomics studies.

## Introduction

Most proteins function through interactions with other proteins to form complexes that carry out a myriad of biological tasks^1,2^. In the past decade, significant effort has been invested in building comprehensive interactome networks for human proteins^3-9^. Many proteins are pleiotropic and carry out diverse functions through interactions with different proteins using distinct interaction interfaces^10^. Also, mutations on the same protein can affect different interactions to cause clinically distinct diseases, depending on the interface where the mutation occurs^11^. Therefore, it is of considerable importance to accurately determine the specific interface that mediates each interaction. However, only about 11K experimentally resolved structures of human protein complexes are reported in the PDB while about 352K human binary protein-protein interactions (PPIs) appear in experimental databases^12^ (Supplementary Fig. 1). Our goal here is to provide reliable predictions of interface residues for the vast majority of human PPIs for which experimental structure information is not available.

Recently, a number of computational methods have demonstrated exceptional capabilities in representation learning, particularly for processing 3D protein structures, leading to much improved performance for protein-protein interface residue prediction^13-16^. Among these is our ensemble deep learning pipeline, PIONEER^16^, designed to generate partner-specific interaction interface residue predictions for experimentally detected PPIs. In contrast, PrePPI^17^ uses structural similarity to predict whether two proteins interact with an efficient scoring function that focuses on interfacial residues^18^. The complementary nature of these two approaches underlines the potential of an integrated framework that combines both. Of note, AlphaFold-based methods^19-21^ have made significant advances in predicting multimeric protein structural models, however, these methods are not yet scalable to model entire interactomes.

Here, we present PIONEER2, an equivariant geometric deep learning pipeline to generate precise partner-specific protein-protein interface residue predictions with high accuracy. PIONEER2 integrates a comprehensive set of biophysical, evolutionary, structural, and sequence features, together with PrePPI-predicted interface information based on structural alignment of query proteins to homologous PDB templates. Extensive benchmark tests show that PIONEER2 is significantly better than PIONEER and other similar algorithms^13-15^. Even compared with the latest AlphaFold3^21^ method, PIONEER2 achieves superior peformance when AlphaFold3 produces low confidence structures. We estimate below that these cases correspond to nearly half of all human PPIs. Together with atomic resolution structures of complexes in the PDB, we established a comprehensive multiscale structurally informed human protein-protein interactome consisting of 352,124 experimentally-detected binary interactions with interface information for each. We then validated PIONEER2 predictions through large-scale mutagenesis experiments which revealed superior performance to AlphaFold3. Furthermore, we found clear concordance between PDB and PIONEER2-predicted interfaces for the frequency of common versus rare population variants as well as significant enrichment of disease mutations. In both cases AlphaFold3 predictions were significantly less successful.

Overall, our results establish PIONEER2 as a powerful tool to accurately predict partner-specific protein-protein interfaces. To further enhance its utility, we provide a database of interfacial residues for all >350k experimentally detected PPIs, which was then combined with disease-associated mutations and their functional annotations to construct the PIONEER2 web server (https://pioneer2.yulab.org) platform as a community resource to advance biomedical studies.

## Results

### An equivariant geometric architecture integrating atomic and residue levels of information

PIONEER2 uses a different design than PIONEER and, specifically, is based on an equivariant geometric deep learning architecture designed to incorporate a comprehensive set of features for predicting protein-protein interfaces. Using features from both interacting partners, PIONEER2 effectively combines biophysical, evolutionary, structural, and sequence information for accurate characterization of interfaces. Importantly, PIONEER2 integrates structural information from 3D spatially neighboring residues.

The architecture is based on the concept of extracting meaningful characteristics from both the target and partner proteins, and then combining these representations to classify whether each residue is part of the interface for the specific interaction. To achieve this, PIONEER2 implements two representation learning branches for sequence and 3D structural information, respectively (Fig. 1a). The sequence branch learns sequence embeddings through a transformer encoder and a CNN encoder. The transformer is appropriate for capturing long-range information, while the CNN is used for local information. Each residue is annotated with biophysical properties, conservation, co-evolution, solvent-accessible surface area (SASA), secondary structure, residue depth, and docking features for an in-depth feature characterization. Compared with our previous models (e.g., PIONEER^16^ and ECLAIR^22^), a key new set of features are derived from the interface information of homologous co-complex structures in the PDB^18^ as predicted by PrePPI^17^. As shown in Fig. 1b, for a given query pair of interacting proteins, PrePPI searches the PDB database with a fast 3D structure alignment algorithm to identify template complexes whose chains are structurally similar (i.e., structural homologs) to the query proteins. Eight different PrePPI scores (see Methods) are assigned to each aligned residue of the query proteins as features in the sequence branch (Fig. 1a). It should be noted that the structure alignment by PrePPI is based solely on protein topology facilitating the detection of remote homologs, which is further enhanced by considering both the full-length query protein and its constituent domains, ensuring optimal coverage and utility of these PrePPI-derived features.

**Fig. 1.**
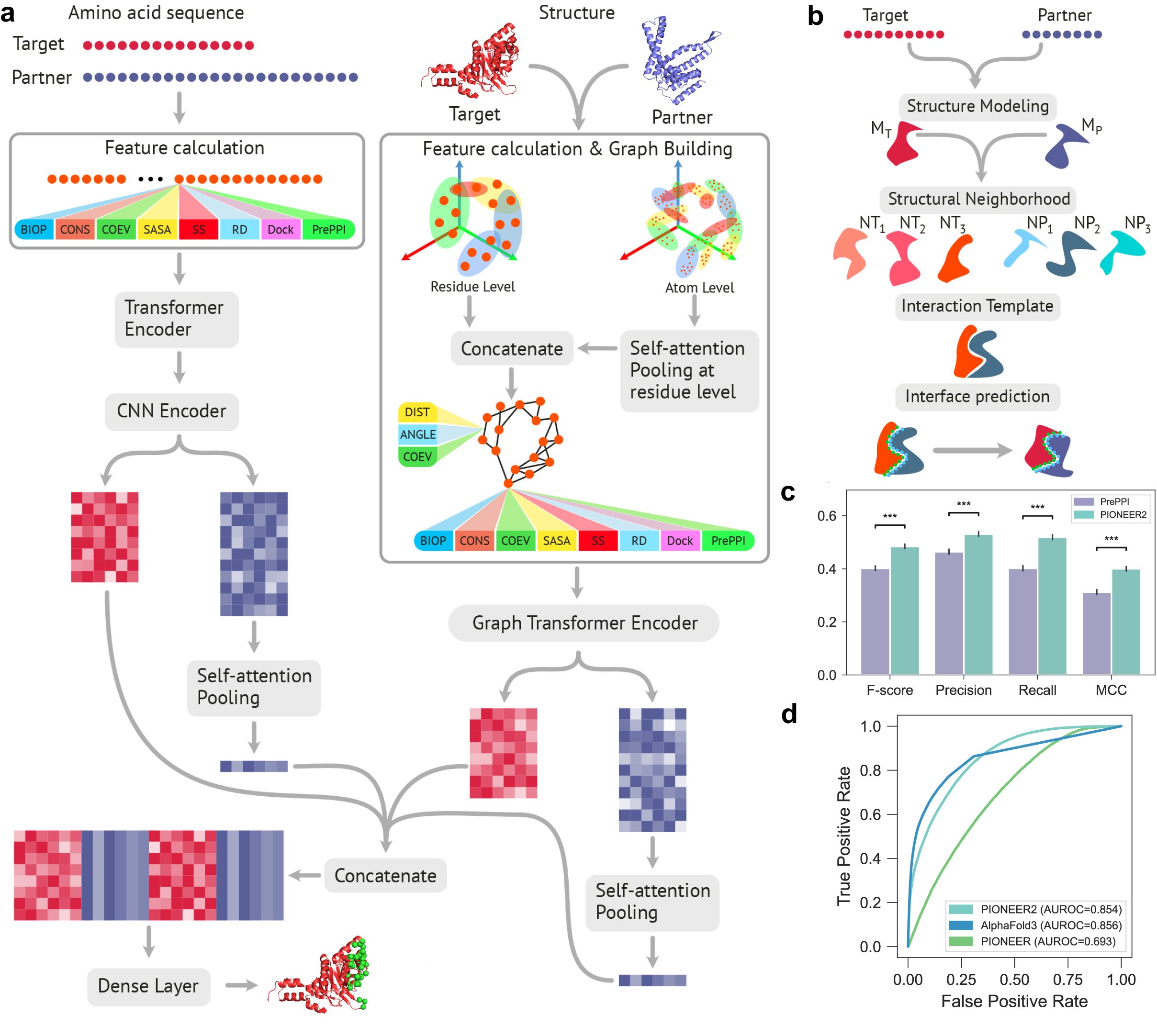
Overview architecture of PIONEER2. **a**, Sequence information for both target and partner proteins is encoded through the Transformer and CNN encoders for the extraction of long-range and short-range dependencies among residues, respectively. The embeddings are then concatenated through self-attention pooling. The structural information of both target and partner proteins is encoded on both residue- and atom-levels, respectively. The residue- and atom-level embeddings are then concatenated through self-attention pooling and are further encoded by the graph Transformer encoders for target and partner, respectively. The resulting embeddings of both target and partner proteins are concatenated through self-attention pooling. Finally, the sequence and structural embeddings are concatenated and are input to the dense layer for the interface residue prediction of the target proteins. **b**, Overview of the PrePPI pipeline which provides features for the sequence branch. Structural models for the query proteins from the AlphaFold Protein Structure Database are structurally superimposed onto chains from the PDB to detect template complexes for modeling. The structure-based sequence alignment of query and template proteins is scored as described in Methods. **c**, Comparison of PrePPI and PIONEER2 performance at recovering the PDB test set. PrePPI interface residues are obtained from the interaction model resulting from structural alignment in panel b. PIONEER2 predictions are based on a probability cutoff that optimizes the F-score. **d**, Comparison of receiver operating characteristic (ROC) curves of PIONEER, PIONEER2 and AlphaFold3 for evaluation on the PDB test set. AlphaFold3 contact probability are used as described in Methods.

In parallel to the sequence branch, the structure branch builds a geometric representation of each protein’s 3D structure at both atomic and residue levels (see Methods). Specifically, atoms and residues are represented in an equivariant manner to preserve geometric properties under rotation and translation. A key step in this process is the integration of information from spatially neighboring atoms and residues—those that are physically close in 3D space, even if they are distant in sequence—allowing the model to capture local environment and long-range dependencies among atoms and among residues. These representations are embedded using a Gaussian mixture model^14^, which captures local spatial distributions by modeling the positions of atoms and residues as a weighted combination of Gaussian components. This enables the model to learn meaningful spatial patterns in the protein 3D structures. Atomic level features are further refined through self-attention pooling and concatenated with residue-level embeddings to form comprehensive structural representations. Every node is annotated with the same features as those in the sequence branch. Additionally, in the structure branch, each edge is annotated with Euclidean distance, angle, and coevolution information. Embeddings are finally processed using a graph transformer encoder.

In both sequence and structural branches, target and partner proteins are represented by a feature matrix, and the partner is further processed by self-attention and concatenated with the target protein. Finally, the embeddings from both sequence and structure branches are integrated into a dense layer, culminating in the final prediction.

### Accurate prediction of protein-protein interfaces enabled by structural homologs

To comprehensively evaluate the performance of PIONEER2, we carefully constructed large labeled datasets with interface information from experimentally determined co-complex structures in the PDB (see Methods) for model training, validation, and testing. Compared with PIONEER labeled sets, we especially prioritize the instances where the same protein interacts with multiple interaction partners using distinct interfaces in our labeled dataset to build a model that better predicts partner-specific interfaces. Furthermore, we ensured that no homologous complexes appear across labeled sets so as to maximize the generalization of our pipeline and to avoid data leakage. In total, the labeled sets contain 4,649, 776 and 802 PDB PPIs for model training, validation and testing, respectively.

As described above, one key addition in PIONEER2 is the binding interface information from structural homologous PDB complexes detected by PrePPI. To mitigate data leakage, for each interaction in the labeled sets, we removed PDB complexes with high sequence homology to both query proteins from the pool of template complexes interrogated by PrePPI. As shown in Fig. 1c, predictive performance on the test set directly using the aligned interface information from the top structural homologs detected by PrePPI is already quite good. Comprehensively incorporating all other features and aggregating neighboring information at both sequence and 3D structure levels in the full PIONEER2 pipeline further significantly improves performance (Fig. 1c).

Since we have previously shown that PIONEER^16^ outperforms recent state-of-the-art interface prediction methods (such as PeSTo^15^, ScanNet^14^, MaSIF-Site^13^, and others), we first compared the performance of PIONEER2 with PIONEER and found that PIONEER2 has significantly improved its predictive performance (Fig. 1d; AUROC increased from 0.69 to 0.85). AlphaFold2^19^ was a breakthrough development in structural modeling for individual proteins and, more recently, many AlphaFold-based methods^23,24^ have been developed to model PPIs. AlphaFold3^21^ is the latest model with state-of-the-art performance for modeling PPIs as well as protein-DNA and -small-molecule interactions. We compared PIONEER2 performance to that of AlphaFold3, using the contact probabilities provided by AlphaFold3 on a per-residue basis (Fig. 1d), and found that PIONEER2 achieves comparable overall performance (AUROC = 0.85, 0.86 for PIONEER2 and AlphaFold3, respectively).

### PIONEER2 significantly outperforms AlphaFold3 in interface residue prediction for nearly half of the human interactome

There are hundreds of thousands of experimentally detected binary PPIs (see Methods) for human proteins already published in the literature, only ∼3.2% of which have structural models in the PDB (Supplementary Fig. 1). It is thus of great interest to predict binding interface information for the large number of human PPIs of unknown structure^12^.

Since AlphaFold3 has become the *de-facto* method for computationally predicted structural models of protein interactions, we carefully analyzed its performance on interactions of known and unknown structure. We generated AlphaFold3 models for the test set of 802 PDB complexes (Fig. 1d) and for 1,024 randomly chosen PPIs of unknown structure from our high-quality literature-curated binary interaction dataset, HINT-HQ-LC-Binary^12^. HINT-HQ-LC-Binary, a subset of HINT-Binary (see below and Methods), contains 44,115 PPIs, each of which has at least two publications using low-throughput detection methods with evidence codes confirming direct, physical binding. Therefore, the HINT-HQ-LC-Binary dataset contains high-quality protein-protein interactions with few false positives^25-27^.

We used the AlphaFold3 ranking score^21^ as the confidence metric for AlphaFold3 models. A higher ranking score indicates a greater likelihood that the complex structure is accurately predicted; ranking scores less than 0 were truncated to 0 for this analysis. The ranking score distributions in Fig. 2a, b illustrate that, overall, AlphaFold3 models for interactions of unknown structure (light blue) are of lower confidence than models for interactions with PDB structures (dark blue): 48% of models for interactions of unknown structure have ranking scores ≤ 0.5 whereas only 23% of models for PDB interactions score that poorly.

**Fig. 2.**
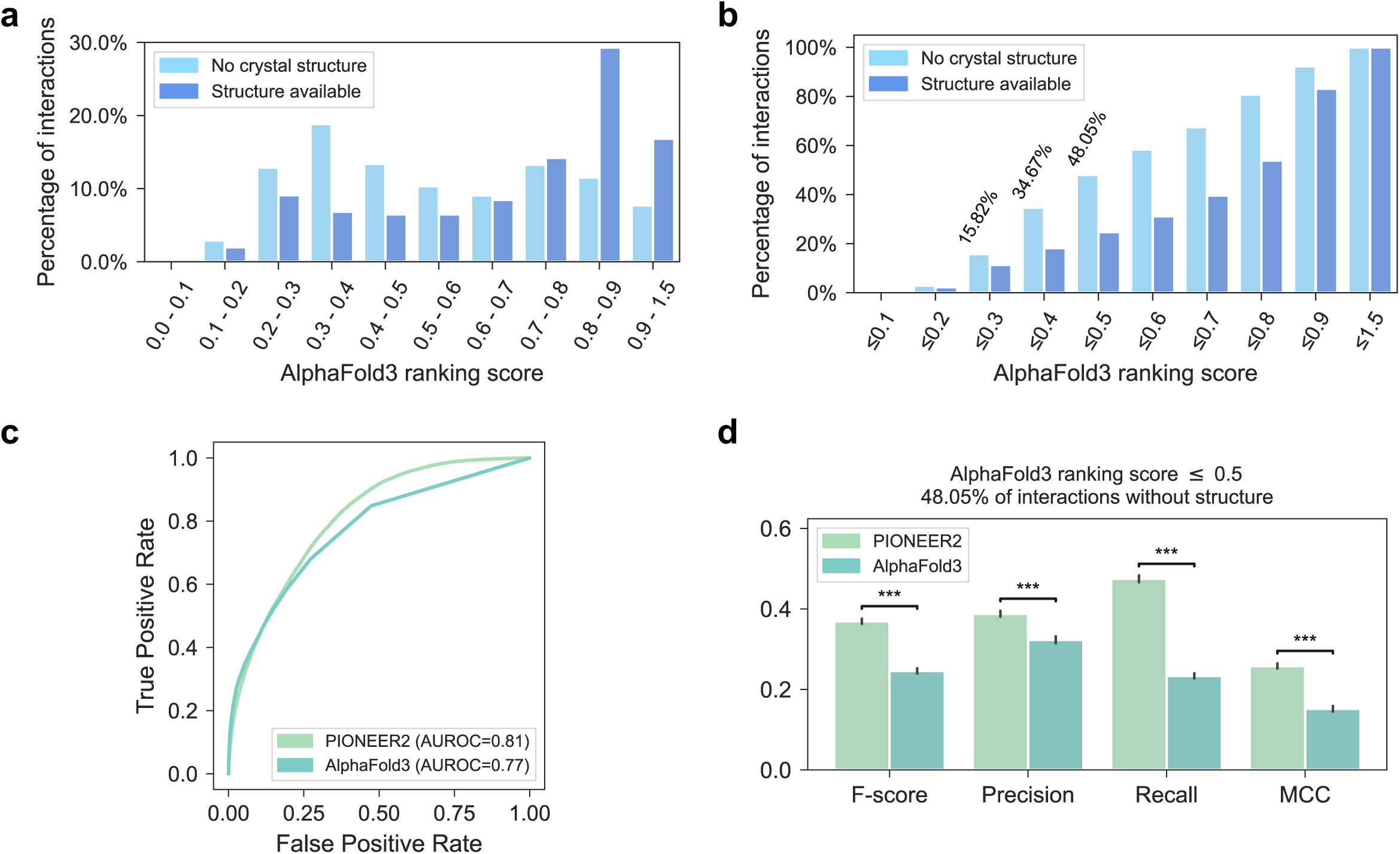
PIONEER2 and AlphaFold3 interface residue predictions for protein interactions of known and unknown structure. **a**, Distribution of AlphaFold3 ranking scores on interactions with and without PDB structures. **b**, Cumulative distribution of AlphaFold3 ranking scores on interactions with and without PDB structures. **c**, Comparison of ROC curves for PIONEER2 and AlphaFold3 using the bootstrapped samples from the test set. AUROC = 0.5 for a random prediction. **d**, Comparison of PIONEER2 and AlphaFold3 on interactions with AlphaFold3 ranking score below 0.5, using the bootstrapped samples. AlphaFold3 predictions are taken from predicted structures with interface residues defined as described in Methods. The vertical axis denotes the metric value. The “***” indicates that the *P*-value is less than 0.001.

Because 97% of the human interactome does not have PDB structures, we sought to evaluate PIONEER2 and AlphaFold3 on PDB structures that have the same distribution of AlphaFold3 ranking scores as the 1,024 interactions of unknown structure (Fig. 2a). To this end we applied a bootstrap resampling strategy to the test set of 802 PDB complexes (see Methods). As shown in Fig. 2c, in this scenario, PIONEER2 (AUROC = 0.81) outperforms AlphaFold3 (AUROC = 0.77). Furthermore, we focused on the more challenging interactions whose AlphaFold3 ranking scores are below 0.5, which accounts for nearly half (48.05%) of all human interactions without known structures (Fig. 2b). Our analysis reveals that PIONEER2 significantly outperforms AlphaFold3 in interface predictions for these interactions, as evaluated by F-score, Precision, Recall, and MCC scores (Fig. 2d).

We then conducted a detailed performance comparison between PIONEER2 and AlphaFold3 across different AlphaFold3 ranking score bins for the 802 PDB PPIs in the test set (without resampling). The results reveal that PIONEER2 consistently and significantly outperforms AlphaFold3 in all bins where the ranking score ≤ 0.5 (Fig. 3 and Supplementary Fig. 2), and achieves comparable performance in bins with higher AlphaFold3 ranking scores (Sup Fig. 2).

**Fig. 3.**
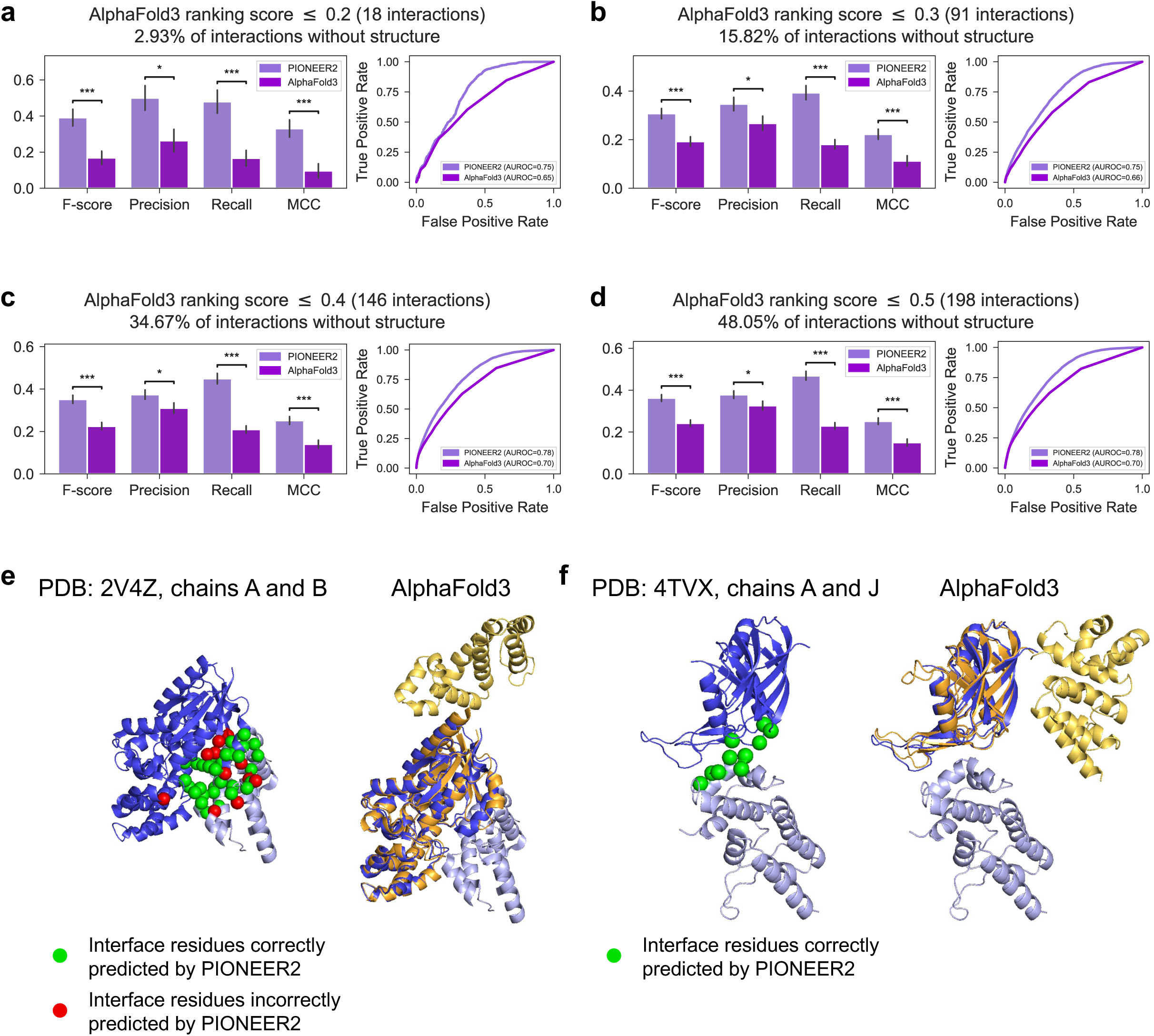
PIONEER2 outperforms AlphaFold3 for interface residue prediction across various AlphaFold3 model confidence levels. **a-d**, Comparison of PIONEER2 and AlphaFold3 on interactions with AlphaFold3 ranking scores below 0.2 (**a**), 0.3 (**b**), 0.4 (**c**) and 0.5 (**d**), respectively. The “*” and “***” indicate that the *P*-value is less than 0.05 and 0.001, respectively. In the bar plots, the vertical axis denotes the metric value. **e-f**, AlphaFold3 predictions on the interactions between the proteins encoded by GNAI3 and RGS2 (PDB: 2V4Z, chains A and B) (**e**) and by casE and casB (PDB: 4TVX, chains A and J) (**f**) demonstrate that AlphaFold3 fails to correctly position the two proteins relative to each other, leading to inaccurate interface predictions. The light blue and dark blue ribbons represent the true co-complex structures, and the yellow and orange ribbons represent the AlphaFold3 predictions. The green and red spheres correspond to correct and incorrect PIONEER2 interface predictions, respectively.

Of course, a key distinction between PIONEER2 and AlphaFold3 is that PIONEER2 focuses exclusively on predicting interface residues, while AlphaFold3 is designed to predict co-complex structures, which is a much more difficult task. Therefore, even minor changes in the relative positions of the two proteins in a structural model can have a significant impact on the accuracy of interface residue predictions by AlphaFold3. Here, we show two examples (Figs. 3e, f) where AlphaFold3 builds good models for the monomers but places them in different orientations relative to the PDB complexes, leading to misidentification of interface residues even when the contact probabilities may be high. Interface residues predicted by PIONEER2 are labeled as green (correct prediction) and red (incorrect) spheres. It is interesting to note that even the technically incorrect predictions (red spheres) by PIONEER2 are in the binding interface area, highlighting the robust performance of PIONEER2, even for cases when AlphaFold3 struggles.

### Construction and validation of a structurally informed human interactome

We compiled HINT-Binary^12^, a comprehensive set of experimentally validated human PPIs which integrates information from commonly used databases, including BioGRID^28^, DIP^29^, IntAct^30^, MINT^31^, iRefWeb^32^, HPRD^33^, PDB^18^, and MIPS^34^, by examining the evidence codes and extent of literature curation for each interaction (see Methods). This procedure yielded 352,124 binary human PPIs, 340,860 of which have no PDB structures. We then used PIONEER2 to predict interfaces for all 340,860 interactions. Since we make a partner-specific interface prediction for every residue in a protein pair, this requires predictions for more than 397 million residue-protein interaction pairs. By combining PIONEER2 interface predictions with the 11,264 PDB interactions, we generated a comprehensive structurally informed human interactome (Fig. 4a), in which all 352,124 interactions have partner-specific interface information at the residue level, together with atomic resolution 3D models for those that appear in the PDB.

**Fig. 4.**
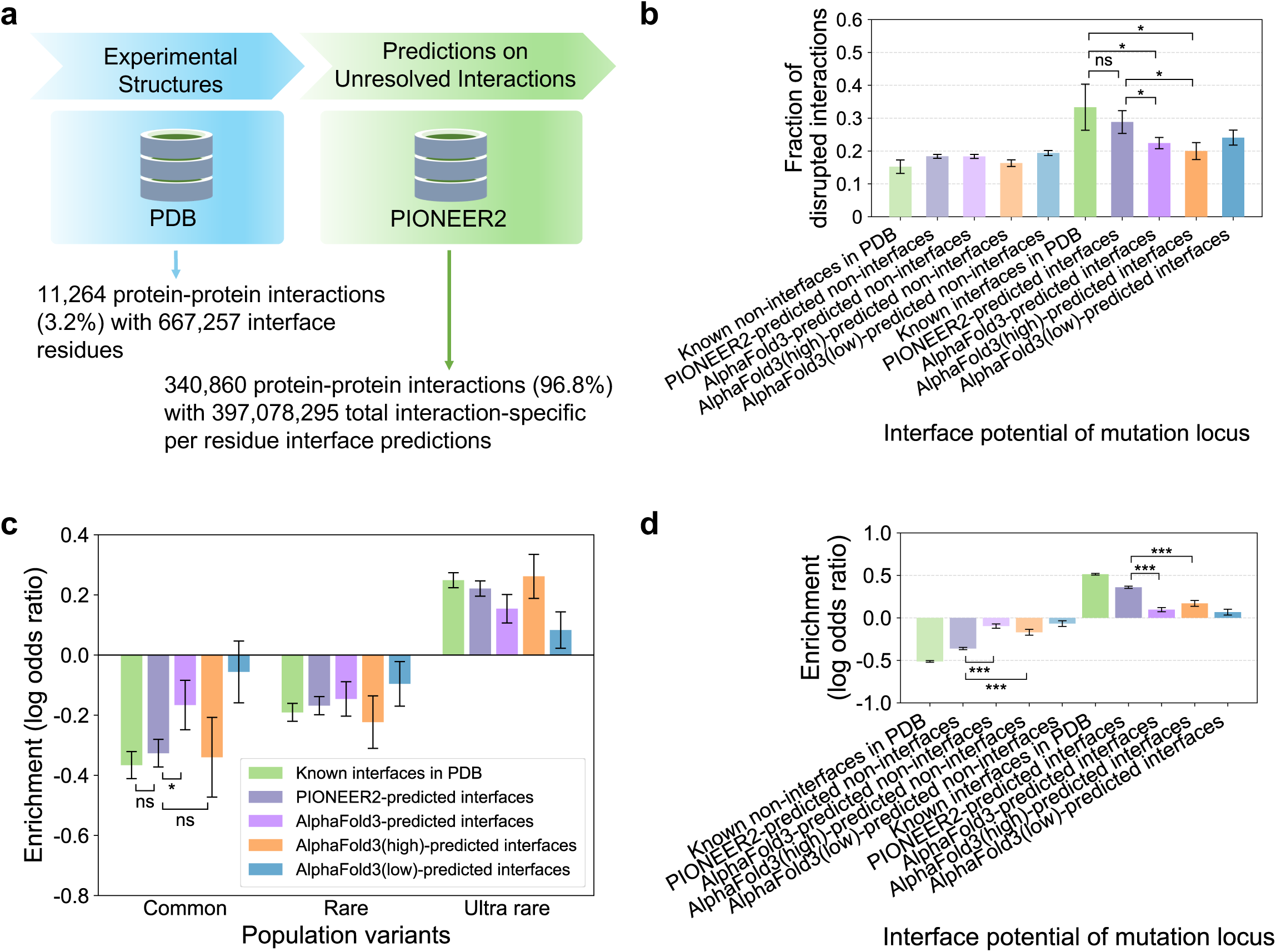
PIONEER2 provides high-quality interface residues for the structurally informed human protein-protein interactome. **a**, The interactome includes interface residues calculated from experimentally determined PDB structures, and PIONEER2 predictions of interface residues for the remaining unresolved interactions. **b**, Fraction of interactions disrupted by random population variants in PDB, PIONEER2-predicted, and AlphaFold3-predicted interface residues. We further divided all AlphaFold3 models into high-quality (“high”: ranking scores > 0.5) and low-quality (“low”: ranking scores ≤ 0.5) subsets. The error bar denotes standard error for the binomial distribution. Significance was determined by the two-sided *z*-test. **c**, Enrichment of population variants in PDB, PIONEER2-predicted, and AlphaFold3-predicted (all, high, and low) interfaces. The error bar denotes standard error for the log odds ratio. **d**, Enrichment of disease-associated mutations in PDB, PIONEER2-predicted, AlphaFold3-predicted (all, high, and low) interfaces. The error bar denotes standard error for the log odds ratio. Significance was determined by the two-sided *z*-test.

We conducted a comprehensive evaluation of the quality of our predicted interface residues by performing large-scale mutagenesis experiments. These experiments measure the fraction of disrupted interactions caused by mutations to our predicted interfacial and non-interfacial residues. We also measured the effects of mutations to known interfacial and non-interfacial residues in PDB co-complex structures. Using our Clone-seq pipeline^35^, we generated 1,866 mutations across 895 proteins and assessed their impact on 5,010 mutation-interaction pairs via a high-throughput yeast-two-hybrid (Y2H) assay. Notably, as shown in Fig. 4b, mutations in both known and PIONEER2-predicted interfacial residues cause disruptions at a much higher rate than those predicted by AlphaFold3, and PIONEER2 significantly outperforms even high-quality AlphaFold3 models (ranking scores > 0.5). Specifically, mutations of non-interfacial residues predicted by PIONEER2 and by AlphaFold3 disrupt PPIs at rates similar to that observed for PDB complexes. In contrast, mutations of PIONEER2-predicted and PDB interfacial residues disrupt interactions at significantly higher rates whereas mutations of AlphaFold3-predicted interfacial residues disrupt interactions at a significantly lower rate, similar to that for non-interfacial residues. These large-scale experiments confirm the high quality of our interface predictions and validate the overall effectiveness of our PIONEER2 pipeline.

We also examined the distribution of population genetic variants and found that their enrichment in PIONEER2-predicted interfaces versus non-interfaces closely aligns with the distributions observed for PDB structures and high-quality AlphaFold3 models (Fig. 4c). The results reveal a significant depletion of common, non-deleterious variants in both known and PIONEER2-predicted, as well as high-quality AlphaFold3-predicted interfaces; interestingly, although the overall AlphaFold3-predicted interface residues exhibit the same behavior, the trend is significantly weaker (Fig. 4c). Furthermore, disease-associated mutations from the Human Gene Mutation Database^36^ (HGMD) are depleted in non-interfaces and enriched in PDB, PIONEER2- and AlphaFold3-predicted interfaces but again, the trend is much weaker in the latter case (Fig. 4d). In fact, the enrichment of disease-associated mutations is significantly higher in PIONEER2-predicted interfaces than that of the high-quality AlphaFold3 models. Beyond providing validation of our predictions, these results establish PIONEER2 as a useful tool in providing a mechanistic explanation of population variants and disease mutations.

### The PIONEER2 3D interactome web server

As a community resource, we developed the PIONEER2 3D interactome web server (https://pioneer2.yulab.org, Fig. 5), which covers all 352,124 experimentally determined binary human PPIs reported in the literature. The PIONEER2 web server also incorporates 161,244 disease-associated mutations across 10,564 disorders cataloged in HGMD^36^ and ClinVar^37^ with precomputed per-disease enrichment analyses on protein interaction interfaces. PIONEER2 offers a user-friendly platform to visualize each protein and its interactors, integrating all available 3D co-complex and monomer structures, and interaction interfaces (mostly predicted by PIONEER2) with comprehensive disease mutation information. This allows users to navigate multiscale, structurally informed interactome networks and pinpoint functionally enriched regions. We anticipate that the PIONEER2 web server will enable the discovery of novel mutation relationships, thereby advancing disease mechanism studies and facilitating personalized treatment development for cancer and other diseases.

**Fig. 5.**
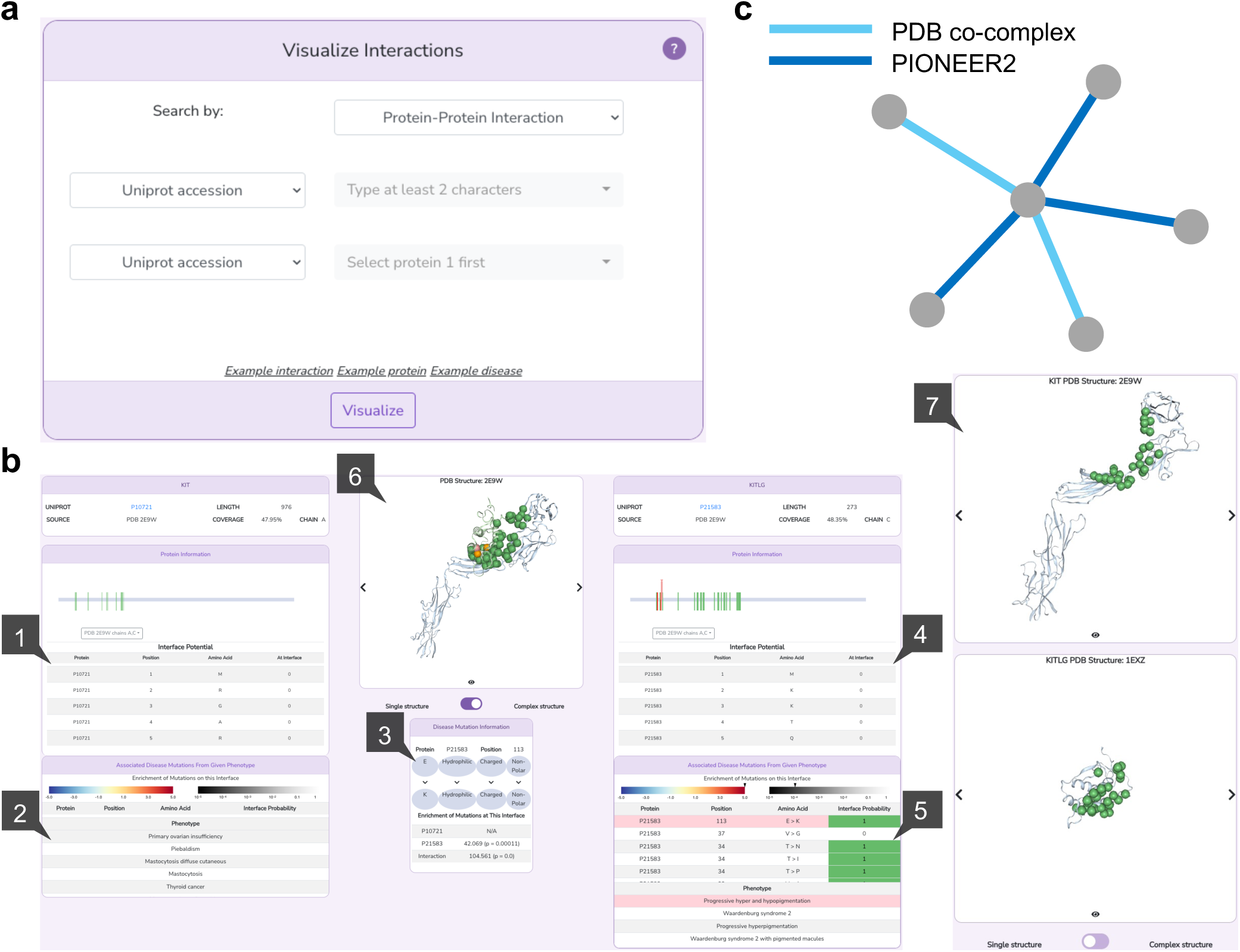
The PIONEER2 3D interactome web server for genome-wide functional exploration. **a**, The PIONEER2 home page, where users can retrieve relevant protein information by searching for a particular interaction (by UniProt ID or gene name), protein, or disease. **b**, The 3D structurally informed PPI and disease-assocated mutation information. Specifically: (1) interface information for each residue on one protein; (2) disease-associated mutation data for that protein; (3) detailed information on mutations associated with a selected disease; (4) interface residue information for each residue on the interacting partner protein; (5) disease-associated mutation data for the partner protein; (6) visualization of the 3D co-complex structure when available, the interface residues and selected disease-associated mutations if available; (7) visualization of individual protein 3D structures and the interface residues. **c**, The visualization of interaction network associated with a specific protein.

## Discussion

In this study, we introduce PIONEER2, an equivariant geometric deep learning framework designed for highly accurate protein-protein interface residue prediction at the whole interactome scale by integrating interface information from PrePPI with a comprehensive set of biophysiochemical, evolutionary, and structural features. Our analysis suggests that AlphaFold-based methods will build lower quality structural models for nearly half of the PPIs of unkown structure in the human interactome compared to those that appear in PDB; for these challenging cases, PIONEER2 outperforms AlphaFold3 in interface residue prediction. Our results thus establish PIONEER2 as a powerful tool for large-scale interface mapping across entire interactomes.

We applied PIONEER2 to the 340,860 human PPIs reported in the literature without experimental structures, which, together with 11,264 PDB PPIs, yields a comprehensive structurally informed human interactome. Using large-scale mutagenesis Y2H experiments, we extensively tested our predictions and found, as shown in Fig. 4b, that PIONEER2-predicted interface residues are validated at the same rate as PDB interface residues. Furthermore, our analyses show that PIONEER2-predicted interface residues have significant enrichment of rare alleles (i.e., functionally important sites) and known disease mutations, to the same degrees as those of PDB interfaces and non-interfaces, confirming the strong biological and functional relevance of PIONEER2 predictions. Of note, AlphaFold3 performance on all three sets of observations is significantly weaker. In summary, our PIONEER2-generated structurally informed human interactome will be an invaluable resource for the study of disease mechanisms and development of personalized treatments.

While PIONEER2 predicts interfacial residues, its performance is significantly enhanced by the inclusion of evidence from PrePPI which is based on a high throughput homology modeling pipeline. However, PIONEER2 performance is superior to that of PrePPI although PrePPI performs quite well on its own. Since, like AlphaFold3, PrePPI provides 3D models of binary PPIs, PIONEER2 predictions can be used as a basis of evaluating and refining PrePPI 3D models by providing more accurate predictions of interfacial residues. The same strategy can of course be applied to AlphaFold3 models. For example, as described above, many of the AlphaFold3 errors likely involve inter-protein orientations rather than the structures of individual monomers. Thus, for low-confidence models, PIONEER2 can be used to assess the quality of the models, and in addition, can potentially be used as the basis for algorithms that further refine or correct protein orientations.

Finally, we expect that PIONEER2 capabilities can be further boosted by implementing advanced architectures that will be developed in the future, and by incorporating novel features, such as sequence language models (e.g., ESM2^38^), and structure language models (e.g., SaProt^39^). With the ongoing progress in sequencing technologies and large-scale genome/exome sequencing projects, such as those from TCGA^40^ and precision medicine initiatives, we envision that PIONEER2’s structurally informed interactome will play an important role in bridging the gap between genomic data and structural proteomics.

## Methods

### Dataset construction

In our HINT^12^ database, we compiled a comprehensive set of experimentally validated binary protein-protein interactions for human, by integrating information from commonly used databases, including BioGRID^28^, DIP^29^, IntAct^30^, MINT^31^, iRefWeb^32^, HPRD^33^, PDB^18^, and MIPS^34^. We carefully examine the evidence codes of each interaction to select only those with experimental evidence to be direct binding. This resulted in 352,124 binary human protein-protein interactions (named “HINT-Binary”), 340,860 of which have no experimental structures; while the other 11,264 have experimentally determined co-complex structures in PDB as of 2024. Among these interactions, we carefully selected a subset (named “HINT-HQ-LC-Binary”) of 44,115 high-quality literature-curated protein-protein interactions that are required to be validated by two separate publications using low-throughput experimental methods.

To construct our labeled sets, partner-specific interface residues were identified from the known co-crystal structures available in PDB. SIFTS^41^ was then used to map the UniProt indices to the PDB indices. The interface residues are determined as residues that are surface residues (≥15% exposed surface) with their relative SASA decreasing by ≥1Å^2^ in the complex. Surface areas were calculated using NACCESS^42^. We processed all available structures in the PDB for each interaction, and considered a residue to belong to the interface if it matches our criterion in at least one of the processed co-crystal structures. Our dataset includes only interactions for which at least 30% of the UniProt residues are covered by co-crystal structures. In addition, to ensure the robustness and generalizability of our models, as well as a fair performance evaluation, we prevented any homologous interactions from being shared between any two datasets that are from the training, validation, and testing splits. Specially, we defined a pair of interactions to be homologous if both proteins in one interaction are homologs to both proteins in the other interaction. We performed three iterations of PSI-BLAST^43^ at an E-value cutoff of 0.001 to determine if two proteins are homologous. This resulted in 4,649, 776, and 802 interactions, corresponding to 2,441,383, 357,276, 326,868 labeled residues for sufficient model training, validation, and testing, respectively (Supplementary Data).

### Graph representation

For a given protein, we created a graph for each residue to enhance structural representations. Each residue is represented by its Cα atom, and the graph includes the 16 closest neighbors to the target residue as nodes with their interactions as edges. Each node and edge are represented by a comprehensive set of features. In specific, in addition to the features used in our previous pipeline, PIONEER^16^ and ECLAIR^22^, which describe each residue in terms of biophysical residue properties, conservation, coevolution, relative SASA, and secondary structure, we have now incorporated three new feature groups for the node representation. These include residue depth, docking metrics, and PrePPI metrics, offering a more comprehensive feature characterization.

We used MSMS^44^ to calculate the residue depth, which describes the average distance of the atoms of a residue from the solvent accessible surface. EquiDock^45^ was used for the rigid body protein-protein docking, after which the interface residues were calculated using the above pipeline. PrePPI^17^ is a pipeline for proteome-wide prediction of known and novel PPIs; PrePPI models and results are available for examination and download from our website. In specific, pairs of proteins, represented by models from the AlphaFold Structure Database^46^, are screened for structural neighbors against a large pool of structures from the PDB database with a fast structure alignment method^47^. For a given pair of query proteins, a suitable interaction template contains structural neighbors of both queries, and the structure-based sequence alignment is examined. The model for the putative PPI is calibrated against the PDB template complex interface, providing several scores that are incorporated in a Bayesian framework to provide a structural modeling likelihood ratio that the query proteins interact The PrePPI feature group includes 8 scores, which are structural modeling likelihood ratio for the overall model, structural similarity between the query protein models and their respective template chain, the number of residue pairs in the template, the number of interacting residue pairs in the template that are preserved in the model, the fraction of interacting residue pairs in the template that are preserved in the model, the number of predicted interfacial residues in the query models that align to interfacial residues in their respective template chains, and the number of interacting residue pairs in the template that are preserved in the model and are predicted to be interfacial residues in the query protein with the program PredUs^48^, respectively.

For the edge, the feature group includes the Euclidean distance, angle and the coevolution between the interacted node pair.

### Model building

To ensure the in-depth capture of structural information, we incorporated the equivariant geometric information at both atomic and residue levels before the processing of the graph Transformer in the structure branch of PIONEER2 architecture. The geometric representation is referred to the literature^14^ to make sure the equivariant properties under rotation and translation. The geometric representations are embedded through Gaussian mixture model at both atomic and residue levels, respectively, and are subsequently concatenated via the self-attention pooling at the amino acid scale. The concatenation is then integrated with the node features along with the edge features for the subsequent graph Transformer embedding.

We compiled a set of representative protein structures from PDB and AlphaFold database for each protein. To enhance the practical applicability of our method more and prevent the memorization of known interfaces, we trained the model using the single protein structures that are not derived from co-crystal or homologous co-complex structures. PDB structures were given the highest priority, while AlphaFold structures were used as a secondary option. The structures are then ranked based on UniProt residue coverage as determined by SIFTS, while ensuring that homologous PDB structures of interacting protein pairs were excluded. For each target protein, residue-level predictions were made using the first corresponding structure that contains the structural information of that residue. For the partner protein, we selected only the structure with the highest UniProt coverage. The PIONEER2 framework was implemented using PyTorch (https://pytorch.org), with the graph Transformer built on the Deep Graph Library (https://www.dgl.ai). The model was trained using binary cross-entropy loss and the Adam optimizer. To accommodate the variable length inputs, we trained the model in a mini-batch mode, processing a single protein pair per batch.

The threshold used by PIONEER2 to define interface residues was determined by optimizing the F-score. AlphaFold3-predicted contact probabilities were used to generate ROC curves for performance evaluation. For the calculation of F-score, precision, recall, and MCC, interface residues in AlphaFold3 structure models are defined as residues that are surface residues (≥15% exposed surface) with their relative SASA decreasing by ≥1Å^2^ in the complex, exactly the same as interface residues defined for PDB structures as described in the “Data Construction” section of the Methods.

### AlphaFold3 predictions for comparison with PIONEER2-generated structurally informed human interactome

For the comparison with AlphaFold3 using mutagenesis Y2H data (Fig. 4b), we generated AlphaFold3-predicted complexes for all 2,159 PPIs that were tested by Y2H. For Fig. 4c,d, we randomly selected 3,806 PPIs from HINT-Binary for AlphaFold prediction to enable the comparison at the interactome level.

### Mutagenesis validation experiments

We conducted mutagenesis experiments by introducing random human population variants from the gnomAD database^49^ into predicted interfaces, known interfaces and non-interfaces. Variants located in predicted interfaces were selected based on PIONEER2 results, while those in known interfaces and non-interfaces from co-crystal structures in PDB served as positive and negative controls, respectively. All selected mutations were introduced using our Clone-seq pipeline^35^. In total, we generated 1,866 mutations across 895 proteins and assessed their effects on 5,010 mutation-interaction pairs—either disrupting or maintaining the interactions—using our high-throughput Y2H assay.

### Y2H assay

We conducted the Y2H assays using the pipeline as previously described^16,22^. We used Gateway LR reactions to transfer all WT/mutant clones into our Y2H pDEST-AD and pDEST-DB vectors. All DB-X and AD-Y plasmids were transformed into the Y2H strains MATα Y8930 and MATa Y8800, respectively. Subsequently, each DB-X MATα transformant (WT and mutants) was individually mated with its corresponding AD-Y MATa transformant (WT and mutants) using automated 96-well procedures, including inoculation of AD-Y and DB-X yeast cultures, mating on YEPD media (incubated overnight at 30 °C) and replica plating onto selective Synthetic Complete media lacking histidine, leucine and tryptophan and supplemented with 1 mM 3-amino-1,2,4-triazole (SC-Leu-Trp-His+3AT), SC-Leu-His+3AT plates containing 1 mg L^−1^ cycloheximide (SC-Leu-His+3AT+CHX), SC-Leu-Trp-Adenine (Ade) plates and SC-Leu-Ade+CHX plates to test for CHX-sensitive expression of the LYS2::GAL1–HIS3 and GAL2–ADE2 reporter genes. We identified spontaneous auto-activators^50^ by growth on plates containing cycloheximide. These plates were incubated overnight at 30 °C and subjected to replica cleaning the following day. The plates were then incubated for an additional three days, after which positive colonies were scored as those growing on SC-Leu-Trp-His+3AT and/or on SC-Leu-Trp-Ade, but not on SC-Leu-His+3AT+CHX or on SC-Leu-Ade+CHX. An interaction was considered disrupted by a mutation if it resulted in a consistent reduction of at least 50% in growth across both reporter genes, relative to the Y2H phenotypes of the corresponding WT allele, as benchmarked by two-fold serial dilution experiments. All Y2H assays were independently repeated three times.

## Data availability

The labeled data used for PIONEER2 model training, validation, and testing are available within the Supplementary Data. The PIONEER2-generated structurally informed human interactome is available at https://pioneer2.yulab.org.

## Code availability

The source code of PIONEER2 is available at https://github.com/haiyuan-yu-lab/PIONEER2.

## Figure legends

**Supplementary Fig. 1. The proportions of the human protein-protein interactome with known (PDB) and unknown structures.**

**Supplementary Fig. 2. Comparison of PIONEER2 and AlphaFold3 across various AlphaFold3 model confidence levels. a-i**, Comparison of PIONEER2 and AlphaFold3 on interactions with AlphaFold3 ranking scores within 0.0-0.2 (**a**), 0.2-0.3 (**b**), 0.3-0.4 (**c**), 0.4-0.5 (**d**), 0.5-0.6 (**e**), 0.6-0.7 (**f**), 0.7-0.8 (**g**), 0.8-0.9 (**h**) and 0.9-1.5 (**i**), respectively. The “*”, “**” and “***” indicate that the *P*-values are less than 0.05, 0.01 and 0.001, respectively. “ns” indicates not significant.

## Acknowledgments

This work was supported by the National Institute of General Medical Sciences (R01GM124559 and RM1GM139738) to HY and (R35GM139585) to B.H, and the Simons Foundation Autism Research Initiative (SFARI-893926) to H.Y.

## Author contributions

H.Y. conceived and oversaw the study; D.X. and H.Y. developed the model; D.M., A.C.N., and B.H. adapted the PrePPI pipeline for model integration; R.F. conducted the Y2H experiments; D.X., M.T., Z.Z., L.L., and H.Y. performed the analyses with discussions with D.M. and B.H.; H.Y. and D.X. wrote the manuscript with input from M.T., D.M., and B.H.; all authors discussed the results and reviewed the manuscript.

