## Supplementary Figures for "Integrating structural homology with deep learning to achieve highly accurate protein-protein interface prediction for the human interactome"

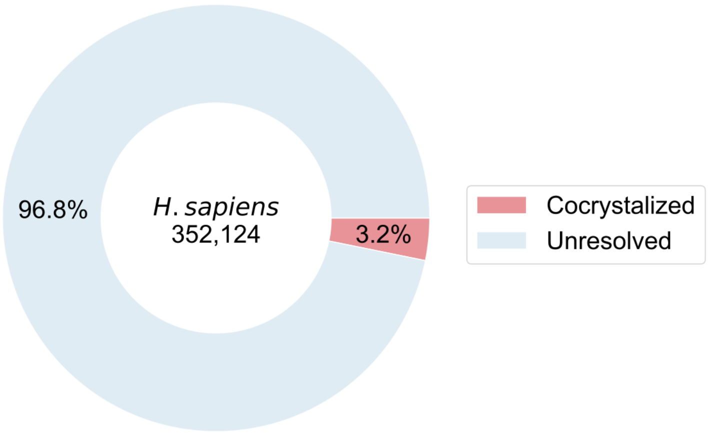


**Supplementary Fig. 1. The proportions of the human protein-protein interactome with known (PDB) and unknown structures.**

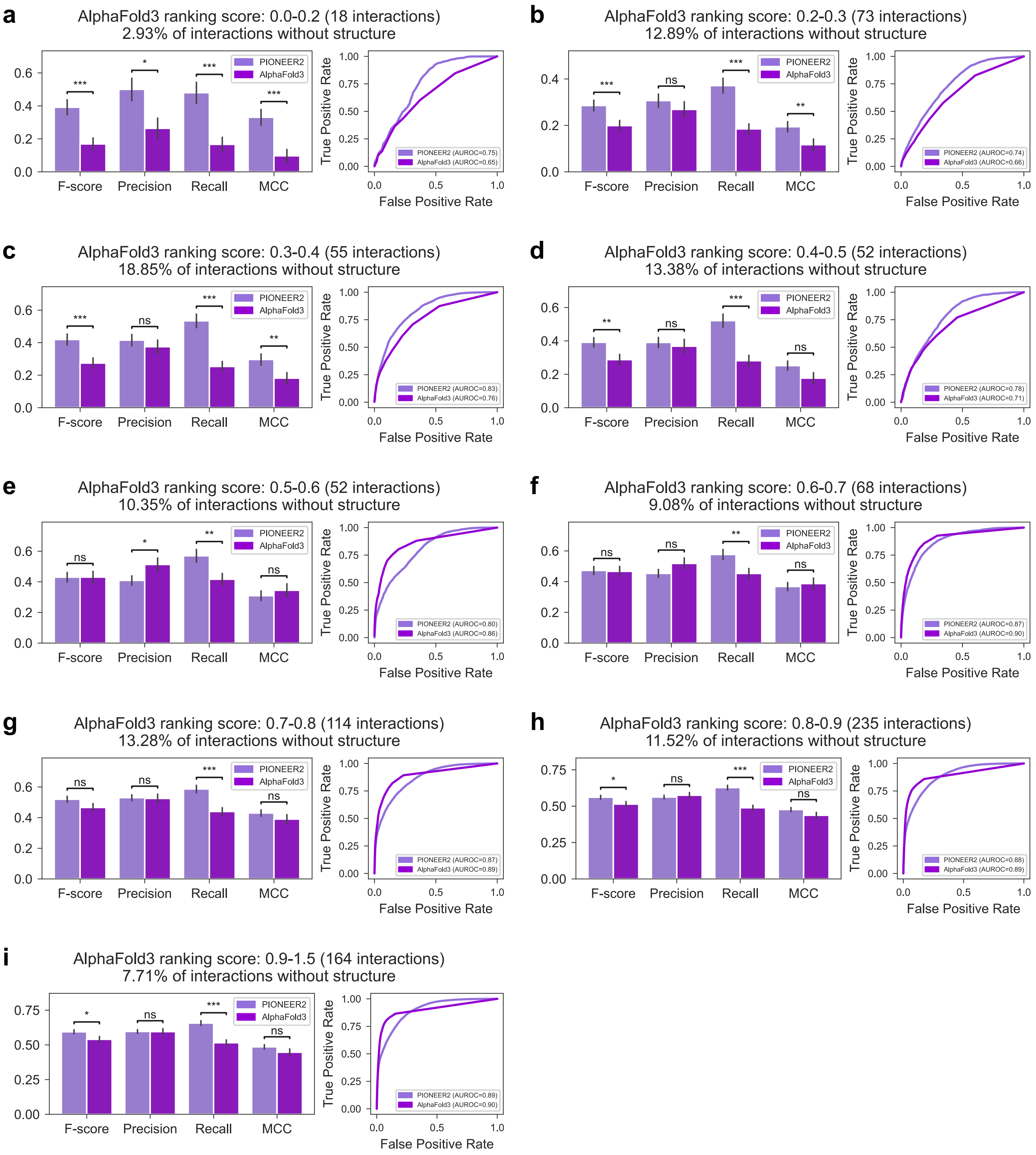


**Supplementary Fig. 2. Comparison of PIONEER2 and AlphaFold3 across various AlphaFold3 model confidence levels.** (**a-i**) Comparison of PIONEER2 and AlphaFold3 on interactions with AlphaFold3 ranking scores within 0.0-0.2 (**a**), 0.2-0.3 (**b**), 0.3-0.4 (**c**), 0.4-0.5 (**d**), 0.5-0.6 (**e**), 0.6-0.7 (**f**), 0.7-0.8 (**g**), 0.8-0.9 (**h**) and 0.9-1.5 (**i**), respectively. The “*”, “**” and “***” indicate that the *P*-values are less than 0.05, 0.01 and 0.001, respectively. “ns” indicates not significant.
